# Angiotensin-converting enzyme governs endogenous opioid signaling and synaptic plasticity in nucleus accumbens

**DOI:** 10.1101/2021.09.08.459469

**Authors:** Brian H. Trieu, Bailey C. Remmers, Carlee Toddes, Dieter D. Brandner, Wei Xie, Swati S. More, Patrick E. Rothwell

## Abstract

Angiotensin-converting enzyme (ACE) regulates blood pressure by cleaving angiotensin I to produce angiotensin II. In the brain, ACE is expressed at uniquely high levels in the striatonigral pathway, but its central function remains poorly understood. We find that ACE degrades an unconventional enkephalin heptapeptide, Met-enkephalin-Arg-Phe, in the nucleus accumbens of mice. ACE inhibition enhanced mu opioid receptor activation by Met-enkephalin-Arg-Phe, causing a cell type-specific long-term depression of glutamate release onto medium spiny projection neurons expressing the Drd1 dopamine receptor. Systemic ACE inhibition was not intrinsically rewarding, but decreased the conditioned place preference caused by fentanyl administration, and enhanced reciprocal social interaction. Our results raise the enticing prospect that central ACE inhibition can boost endogenous opioid signaling for clinical benefit, while mitigating risk of addiction.

## Main Text

As the underlying mechanisms of brain disorders are defined with ever-increasing precision, there is a growing need for interventions that specifically target dysfunctional elements of brain circuits (*1*). A variety of brain conditions, ranging from addiction (*2*) to autism (*3*) to chronic pain (*4*), involve imbalanced output of nucleus accumbens (NAc) medium spiny projection neurons expressing dopamine receptor Drd1 (D1-MSNs) or Drd2 (D2-MSNs). This imbalance has proven difficult to correct with standard interventions, because these two MSN subtypes are physically intermingled, receive synaptic inputs from common sources, and have similar molecular profiles. A rare exception is angiotensin-converting enzyme (ACE), a dipeptidyl carboxypeptidase expressed at uniquely high levels in the striatonigral pathway formed by D1-MSNs (*5*), providing a more precise therapeutic target than dopamine receptors that have widespread expression.

ACE inhibitors have been used for decades as a safe and efficacious treatment for hypertension. Patients taking centrally active ACE inhibitors can experience relief from depression and improved quality of life (*6-13*), though the mechanism of action is unclear. Rodents that exhibit depression-like behavior after chronic stress have upregulated ACE expression in the NAc (*14*), but it is not known if ACE operates canonically in the NAc by converting angiotensin I to angiotensin II (Fig. 1A). In rodents, centrally active ACE inhibitors produce antidepressant-like effects that can be blocked by opioid receptor antagonists (*15-17*), instead suggesting ACE may cleave and degrade a peptide ligand for opioid receptors (Fig. 1B).

**Fig. 1.**
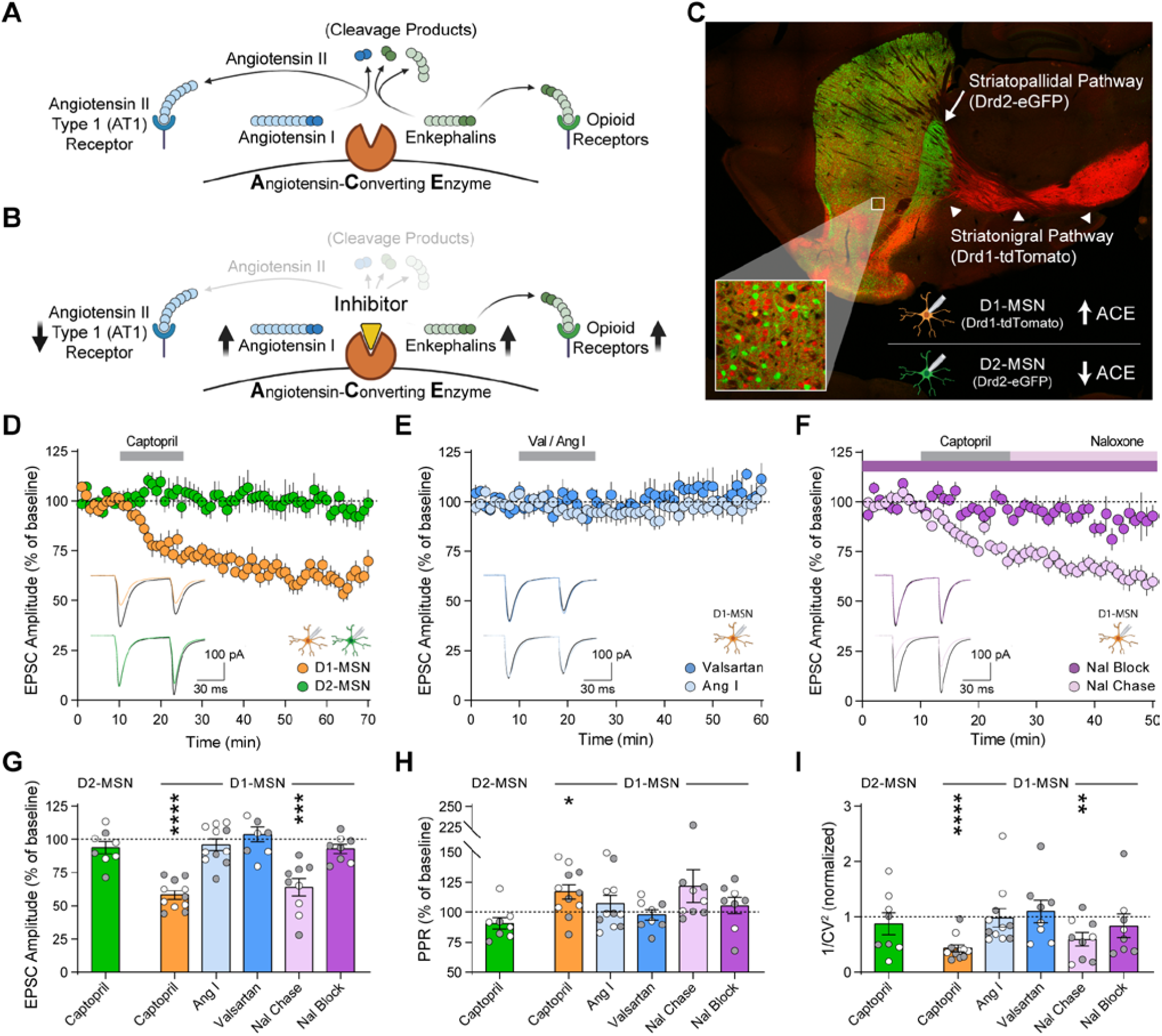
ACE inhibition reduces excitatory input onto D1-MSNs via endogenous opioid signaling. (**A**, **B**) Schematic of canonical angiotensin and non-canonical enkephalin regulation by ACE (A), and the effects of ACE inhibition by captopril (B). (**C**) Drd1-tdTomato expression (red) in D1-MSNs that form the striatonigral pathway (arrowheads) and express ACE, as well as Drd2-eGFP expression (green) in D2-MSNs that form the striatopallidal pathway (arrow). (**D-F**) EPSC amplitude before, during, and after 15 min bath perfusion (grey bar) of captopril (10 µM) in D1-MSNs (orange, *n*=11) or D2-MSNs (green, *n*=8) (D); AT1 receptor antagonist valsartan (2 µM, dark blue, *n*=8) or angiotensin I peptide (1 µM, light blue, *n*=11) in D1-MSNs (E); or captopril (10 µM) in continual presence of opioid receptor antagonist naloxone (10 µM, dark purple, *n*=8) or chased by naloxone (10 µM, light purple, *n*=9) in D1-MSNs (F). Bottom-left insets show representative traces before (black lines) and after bath application (last 5 min of recording, colored lines). (**G-I**) EPSC parameters averaged in the last 5 min of each recording, expressed as percentage of baseline prior to drug application: EPSC amplitude (G) paired-pulse ratio (H), and inverse of squared-coefficient of variation (I). Data are mean ± s.e.m. for all panels; open and closed circles indicate recordings from female and male mice, respectively. \**P*<0.05, \*\**P*<0.01, \*\*\**P*<0.001, \*\*\*\**P*<0.0001, ANOVA followed by one-sample *t*-test versus baseline; see Table S1 for complete statistics.

In the striatum and other brain regions, inhibitors of ACE and other peptidases can be combined to regulate excitatory synaptic transmission in an opioid-dependent fashion (*18, 19*). To measure functional effects of ACE inhibition on NAc excitatory synaptic transmission, we performed whole-cell patch-clamp recordings from MSNs in acute brain slices. To differentiate recordings from D1-MSNs and D2-MSNs, we used double-transgenic Drd1-tdTomato/Drd2-eGFP reporter mice (Fig. 1C). Brief exposure to captopril (10 µM), a prototypical ACE inhibitor (*20*), caused long-term depression (captopril-LTD) of excitatory synaptic transmission onto D1-MSNs, with no effect on D2-MSNs (Fig. 1D).

By blocking the conversion of angiotensin I to angiotensin II, captopril and other ACE inhibitors canonically prevent activation of the angiotensin II type 1 (AT1) receptor, and increase levels of angiotensin I (Fig. 1B). However, LTD was not observed in D1-MSNs when slices were exposed to valsartan (2 µM), an AT1 receptor antagonist, or exogenous angiotensin I peptide (1 µM) (Fig. 1E). In contrast, captopril-LTD in D1-MSNs was blocked in the continuous presence of naloxone (10 µM), an opioid receptor antagonist, but was not reversed by chasing captopril with naloxone (Fig. 1F). This demonstrates that captopril induces long-term synaptic changes via transient opioid receptor activation. Captopril-LTD in D1-MSNs was associated with an increase in paired-pulse ratio and a decrease in 1/CV^2^ (Fig. 1, G to I), consistent with a decrease in glutamate release caused by activation of presynaptic opioid receptors (*18, 21*). ACE thus appears to regulate NAc synaptic transmission by a non-canonical mechanism that may involve degradation of endogenous opioid peptides, which is blocked by captopril to enhance endogenous opioid signaling (Fig. 1B).

Local release of enkephalin peptides by D2-MSNs can regulate excitatory synaptic input to D1-MSNs (*22*). While ACE can cleave enkephalin peptides, it is not principally responsible for degradation of conventional Met-enkephalin (Tyr-Gly-Gly-Phe-Met) or Leu-enkephalin (Tyr-Gly-Gly-Phe-Leu) in brain tissue (*23, 24*). In addition to these conventional enkephalin pentapeptides, the proenkephalin gene (*Penk*) also encodes Met-enkephalin-Arg-Phe (Tyr-Gly-Gly-Phe-Met-Arg-Phe, “MERF”), a heptapeptide expressed at high levels in the NAc (*25-28*). Although often viewed as a precursor to Met-enkephalin, MERF itself has high binding affinity for opioid receptors (*29*), is more potent than conventional Met-enkephalin (*30*), and can be degraded by ACE (*31-33*). Using liquid chromatography-tandem mass spectrometry, we developed an assay to simultaneously quantify extracellular levels of these enkephalin congeners and other neuropeptides released from intact mouse brain slices (Fig. 2A and fig. S1). After stimulation with a high concentration of KCl (50 mM) to evoke neuropeptide release, we observed increased extracellular levels of MERF as well as Met-enkephalin and Leu-enkephalin, along with dynorphins and substance P (Fig. 2B). Concentrations of MERF released from isolated NAc tissue punches were higher than dorsal striatum tissue punches (fig. S2, A to C). We could not detect appreciable levels of angiotensin II, further supporting a non-canonical function for ACE in the NAc. Signals corresponding to MERF, Metenkephalin, and Leu-enkephalin were absent in constitutive *Penk* knockout mice, while signals corresponding to other neuropeptides were preserved (fig. S2, D to I), validating the specificity and sensitivity of our approach.

**Fig. 2.**
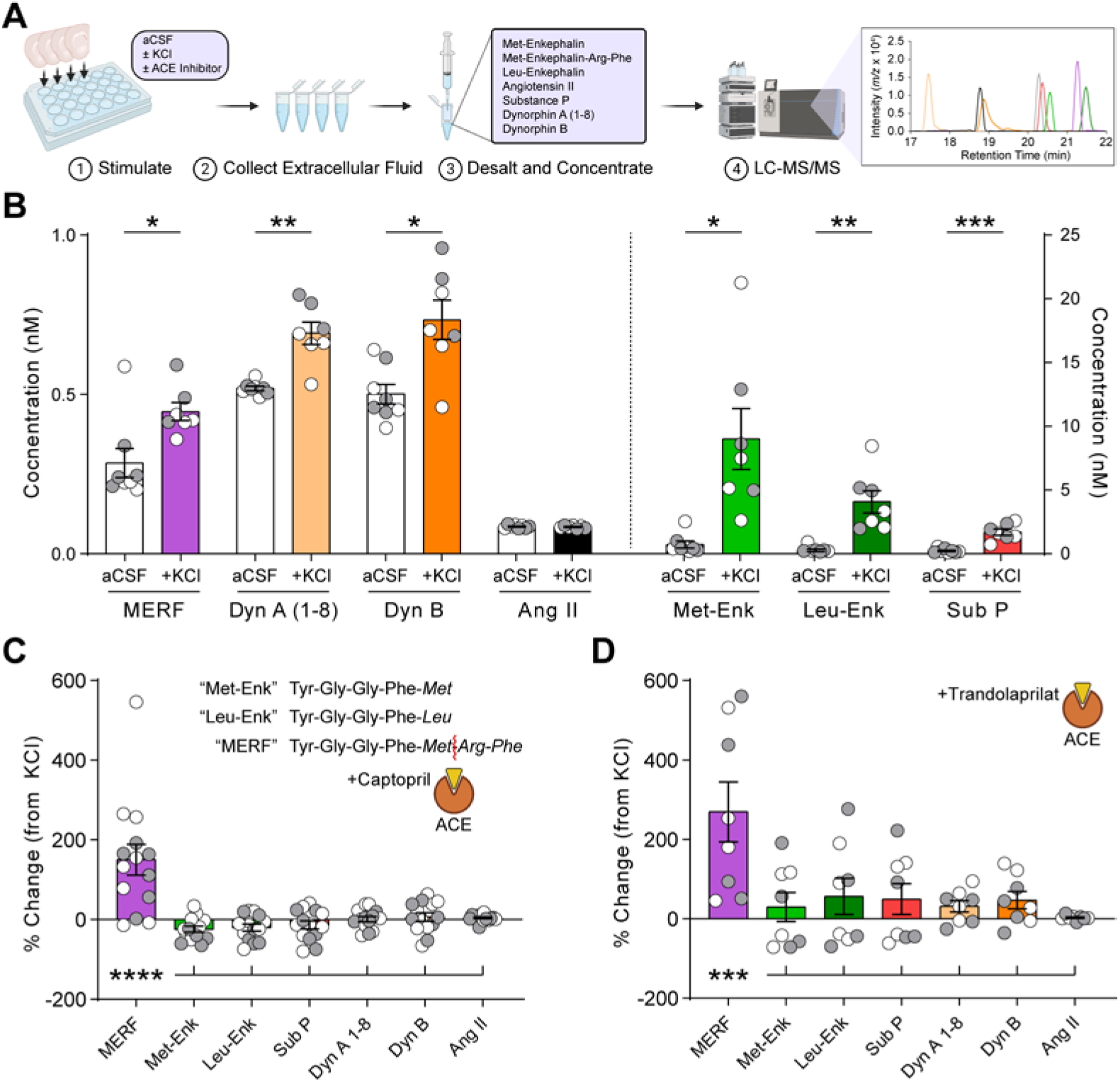
ACE selectively degrades MERF in the extracellular space. (**A**) Schematic of neuropeptide quantification from coronal brain slices using LC-MS/MS. (**B**) Extracellular neuropeptide levels from slices submerged in normal aCSF or a solution containing high KCl (50 mM). (**C, D**) Change in extracellular neuropeptide concentrations after incubation in captopril (10 µM) (C) or trandolaprilat (1 µM) (D), relative to KCl stimulation without ACE inhibitors. Inset shows enkephalin amino acid sequences and site of enzymatic cleavage of MERF by ACE (red line). Data are mean ± s.e.m. for all panels; open and closed circles indicate samples from female and male mice, respectively. \**P*<0.05, \*\**P*<0.01, \*\*\**P*<0.001, \*\*\*\**P*<0.0001, ANOVA followed by simple effect test (B) and ANOVA followed by Fisher’s LSD post-hoc test (C, D); see Table S1 for complete statistics.

The *Penk* gene contains a single copy of both MERF and Leu-enkephalin (*26, 27*), but evoked release of MERF was lower than Leu-enkephalin, suggesting MERF may be rapidly degraded in the extracellular space. The addition of captopril (10 µM) to inhibit ACE robustly increased extracellular levels of MERF, but had minimal impact on conventional enkephalins or other neuropeptides (Fig. 2C). We observed a similar pattern using trandolaprilat, a newer lipophilic ACE inhibitor with increased specificity and potency (Fig. 2D). In contrast, extracellular levels of MERF were not affected by pharmacological inhibition of aminopeptidase N and neprilysin, the enzymes responsible for degrading conventional enkephalins (*34*) (fig. S3A). A cocktail of inhibitors for all three enzymes blocked degradation of enkephalins as well as other neuropeptides (fig. S3B). These data reveal a double-dissociation in the degradation of conventional enkephalins by aminopeptidase N and neprilysin, versus the degradation of MERF by ACE.

To investigate how MERF regulates NAc synaptic transmission, we measured miniature excitatory postsynaptic currents (mEPSCs). Increasing concentrations of MERF caused a dose-dependent decrease in mEPSC frequency, without altering mEPSC amplitude (Fig. 3, A and B), consistent with a presynaptic reduction of glutamate release probability. Both MERF and Met-enkephalin (fig. S4, A to C) had similar effects on D1-MSNs and D2-MSNs, suggesting presynaptic terminals onto both cell types are equally sensitive to endogenous opioids. We used these data to construct dose-response curves, and found that MERF (IC_50_ = 438.4 nM; Fig. 3C) was more potent than Met-enkephalin (IC_50_ = 993.3 nM; fig. S4D), as previously described for analgesic activity (*30*).

**Fig. 3.**
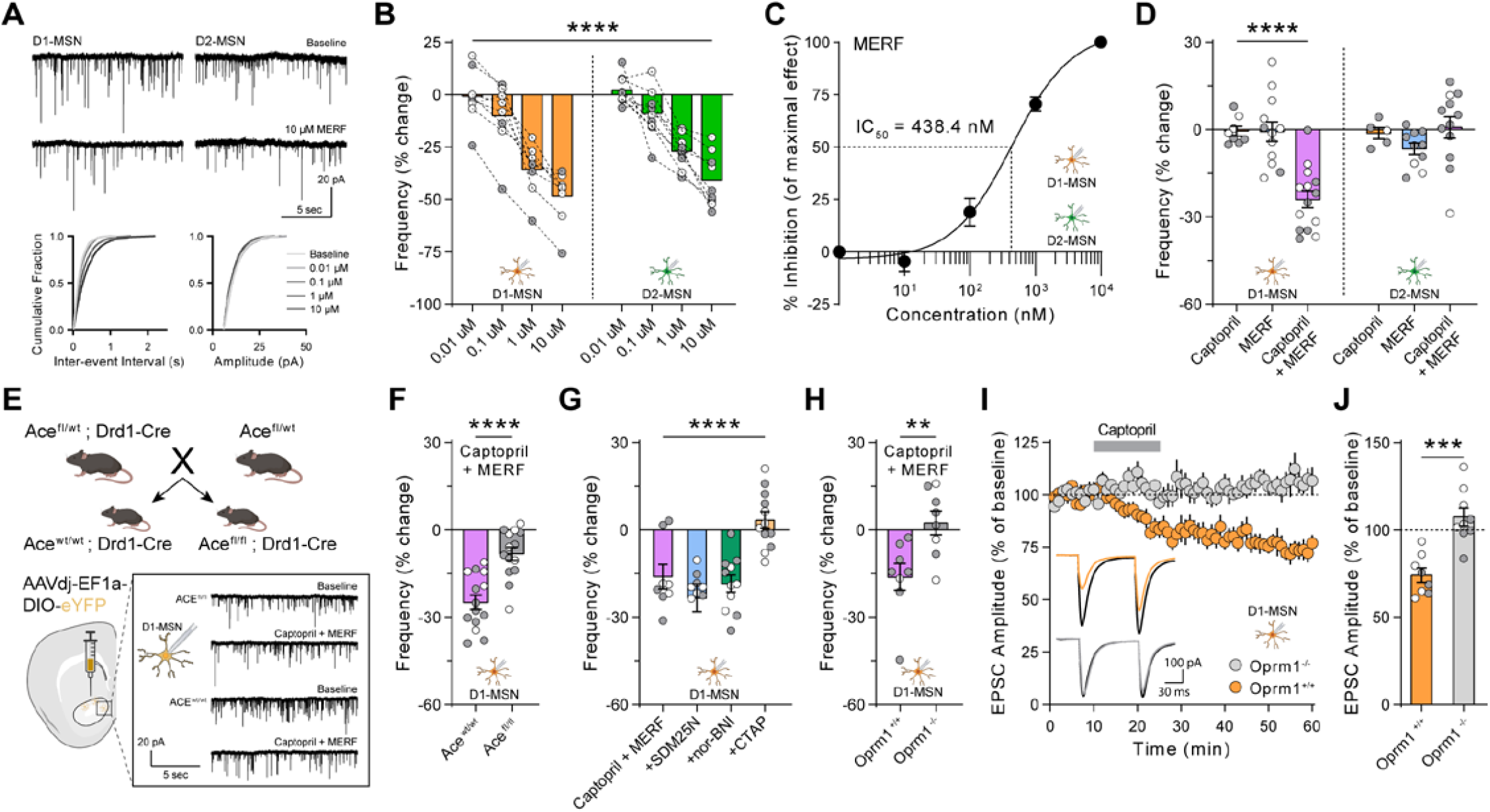
MERF and captopril synergistically depress glutamate release via presynaptic mu opioid receptors. (**A**) Top, representative traces of mEPSCs from D1-MSNs (left) and D2-MSNs (right) during baseline and after bath perfusion of MERF (10 µM). Bottom, cumulative fraction plots of inter-event interval (left) and amplitude (right) of mEPSC events at increasing MERF concentrations (0.01 µM – 10 µM). (**B**) MERF caused a dose-dependent decrease in mEPSC frequency in D1-MSNs (left, orange, *n*=8) and D2-MSNs (right, green, *n*=9). (**C**) Sigmoidal interpolation of MERF dose-response normalized to maximal frequency change at 10 µM (IC_50_: 438.4 nM, 95% CI: 278.7 – 689.7 nM, *n*=17). (**D**) mEPSC frequency after combined captopril (10 µM) and/or threshold MERF (100 nM) in D1-MSNs (left, *n*=14) and D2-MSNs (right, *n*=12). (**E**) Top, genetic cross to obtain control (Ace^wt/wt^; Drd1-Cre) and conditional ACE knockout mice (Ace^fl/fl^; Drd1-Cre). Bottom, stereotaxic injection of Cre-dependent eYFP reporter virus to identify NAc D1-MSNs for whole-cell recordings. Bottom-right inset shows representative traces from control and ACE knockout mice. (**F**) Combined effect of captopril and MERF (100 nM) after conditional deletion of ACE from D1-MSNs (grey, *n*=14) versus control (purple, *n*=14). (**G**) Combined effect of captopril and MERF (100 nM) in the presence of selective antagonists of mu (CTAP, 1 µM, orange, *n*=12), delta (SDM25N, 0.5 µM, blue, *n*=9), or kappa (nor-BNI, 0.1 µM, green, *n*=11) opioid receptors. (**H**) Combined effect of captopril and MERF (100 nM) in Oprm1^-/-^ knockout mice (grey, *n*=8) and Oprm1^+/+^ littermates (purple, *n*=8). (**I**, **J**) EPSC amplitude time course (I) or average during last 5 min (J) of Captopril-LTD (10 µM) in Oprm1^+/+^ (*orange*, n=8) and Oprm1^-/-^ mice (grey, *n*=9). Inset shows representative traces before captopril (black lines) and during last 5 min (color lines). Data are mean ± s.e.m. for all panels; open and closed circles indicate recordings from female and male mice, respectively. \**P*<0.05, \*\**P*<0.01, \*\*\**P*<0.001, \*\*\*\**P*<0.0001, ANOVA concentration main effect (B), ANOVA treatment simple effect in D1-MSNs (D), ANOVA genotype or treatment main effect (F, G, J), and two-sample *t*-test (H); see Table S1 for complete statistics.

These experiments identified a threshold MERF concentration (100 nM) that did not reliably affect synaptic transmission. Captopril alone (10 µM) also had no effect on frequency or amplitude of mEPSCs, which are measured in the absence of stimulation normally required for release of endogenous opioids (*18, 22*). However, the combination of captopril and a threshold concentration of MERF caused synergistic depression of mEPSC frequency in D1-MSNs, but not D2-MSNs (Fig. 3D and fig. S5). Inhibitors of aminopeptidase N and neprilysin did not enhance the effects of MERF, but did potentiate the effects of a threshold concentration of Metenkephalin (100 nM) in both D1-MSNs and D2-MSNs (fig. S4, E to I). These data provide corroborating evidence that MERF is specifically degraded by ACE in the NAc, and indicate MERF is not merely a precursor to Metenkephalin (*32*).

Presynaptic terminals onto both D1-MSNs and D2-MSNs were equally sensitive to endogenous opioids, suggesting postsynaptic expression of ACE makes D1-MSNs uniquely sensitive to the effects of captopril. To test this hypothesis, we crossed floxed ACE mice with Drd1-Cre mice, generating littermate offspring carrying Drd1-Cre that were either homozygous floxed (“Ace^fl/fl^”) or control (“Ace^wt/wt^”) (Fig. 3E). Quantitative RT-PCR on striatal tissue punches from Ace^fl/fl^ mice showed a large reduction of ACE mRNA expression relative to control mice (fig. S6A), confirming that D1-MSNs represent a primary source of ACE expression. To identify D1-MSNs for whole-cell patch-clamp recordings, all mice received stereotaxic injection of a Cre-dependent eYFP reporter virus into the NAc (Fig. 3E). Combined application of captopril (10 µM) and threshold MERF (100 nM) significantly depressed mEPSC frequency in D1-MSNs from Ace^wt/wt^ control mice, but this effect was absent from Ace^fl/fl^ mice lacking ACE expression in D1-MSNs, with no change in mEPSC amplitude in either group (Fig. 3F and fig. S6, B to D). These data confirm that postsynaptic expression of ACE by D1-MSNs is specifically necessary for the enhancement of MERF signaling by captopril.

To determine the opioid receptor subtype engaged by ACE inhibition, we recorded mEPSCs in D1-MSNs and applied captopril (10 µM) with threshold MERF (100 nM), in the presence of selective opioid receptor antagonists. Blocking delta opioid receptors with SDM25N (500 nM) or kappa opioid receptors with nor-BNI (100 nM) did not prevent the decrease in mEPSC frequency, but this effect was completely blocked by the mu opioid receptor antagonist CTAP (1 µM), with no change in mEPSC amplitude (Fig. 3G and fig. S7, A to D). To confirm a specific role for mu opioid receptors, we crossed Drd1-tdTomato reporter mice with constitutive mu opioid receptor (*Oprm1*) knockout mice, generating offspring lacking functional mu opioid receptors (Oprm1^-/-^) as well as littermate controls (Oprm1^+/+^) (fig. S7E). The synergistic effect of captopril (10 µM) and threshold MERF (100 nM) on mEPSC frequency in D1-MSNs was absent from Oprm1^-/-^ mice (Fig. 3H and fig. S7, F to I). Captopril-LTD of evoked EPSCs in D1-MSNs was also absent from Oprm1^-/-^ mice (Fig. 3, I to J). These results demonstrate a high binding affinity of MERF for the mu opioid receptor, which is unique compared to the higher binding affinity of conventional enkephalins for the delta opioid receptor (*29*).

We next evaluated the potential of ACE inhibition to harness cell type-specific regulation of endogenous opioid signaling in behaving mice following systemic administration of captopril. The strengthening of excitatory synaptic input to D1-MSNs and activation of these cells promotes the rewarding effects of addictive drugs and drug-seeking behavior (*35-38*). We investigated the rewarding properties of fentanyl, a potent synthetic opioid, using an unbiased place conditioning assay (Fig. 4A). Mice exhibited robust conditioned place preference (CPP) for a context paired with fentanyl (0.04 mg/kg, s.c.), but the magnitude of CPP was significantly attenuated when captopril (30 mg/kg, i.p.) was injected prior to fentanyl (Fig. 4, B and C), with no effect on fentanyl-induced locomotor stimulation during conditioning (fig. S8, A and B). Captopril itself was not rewarding or aversive in the place conditioning assay (Fig. 4,D to F) and did not alter locomotion (fig. S8, C and D). In a test of social interaction between two freely moving mice, captopril administration increased the amount of social interaction (Fig. 4, G to K), ruling out a general disruption of motivated behavior. Instead, the increase in social interaction following captopril administration is consistent with enhanced mu opioid signaling in the NAc (*39*). The attenuation of fentanyl reward by captopril is consistent with the selective reduction of the excitatory synaptic drive to D1-MSNs, as shown in our whole-cell patch clamp recordings (Fig. 4L). Together, our data illustrate the utility of central ACE inhibition to selectively modulate the activity of D1-MSNs, via regulation of endogenous opioid signaling (Fig. 4M).

**Fig. 4.**
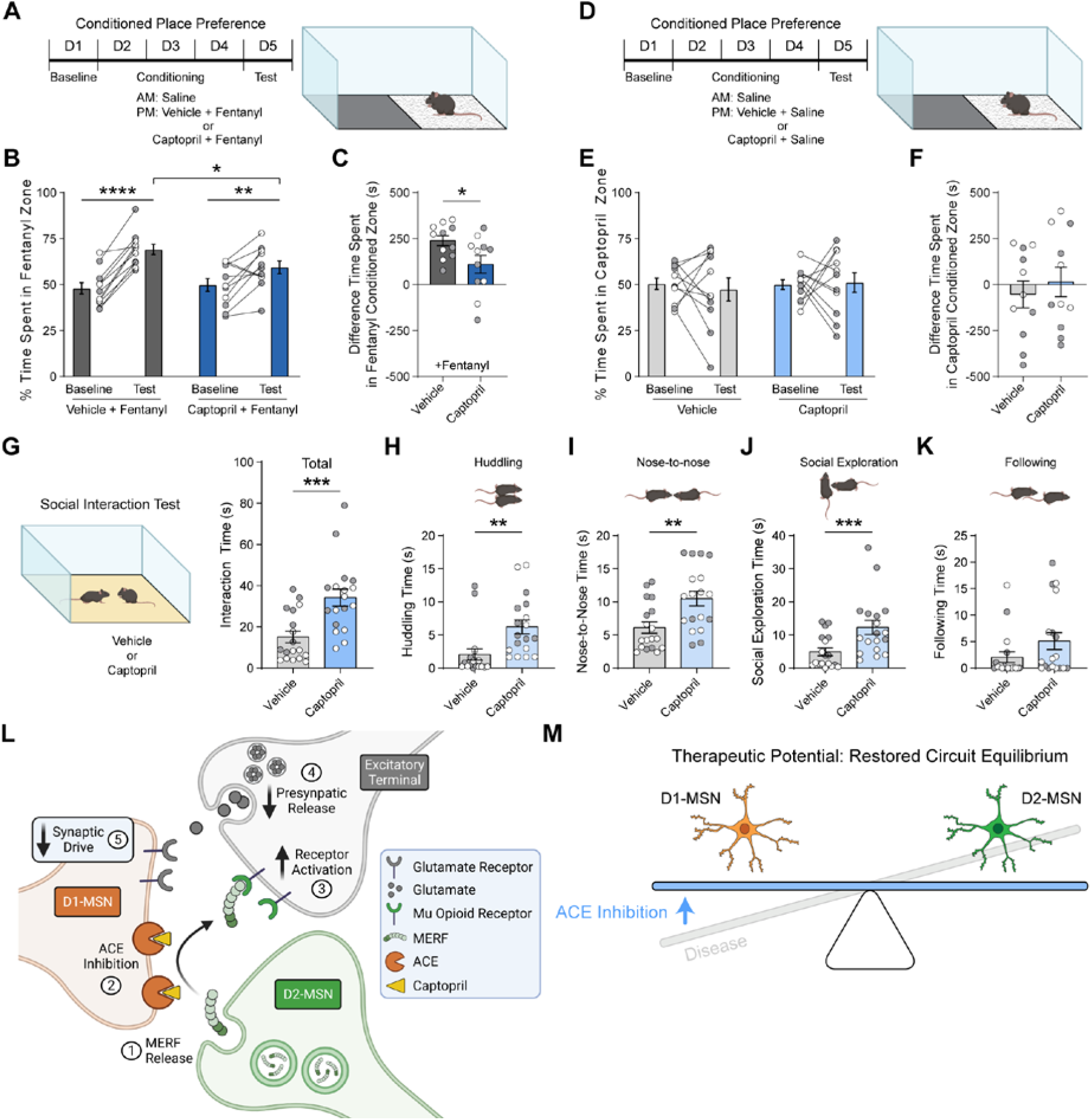
Systemic captopril reduces fentanyl reward and increases sociability. (**A**-**C**) Schematic of unbiased place conditioning procedure (A), with percent time on fentanyl side (B) and CPP score (C) for groups receiving fentanyl (0.04 mg/kg, s.c.) preceded by vehicle (*n*=11, dark grey) or captopril (30 mg/kg, i.p.; *n*=11, dark blue). (**D**-**F**) Schematic of unbiased place conditioning procedure (D), with percent time on fentanyl side (E) and CPP score (F) for groups receiving saline preceded by vehicle (*n*=11, grey) or captopril (30 mg/kg, i.p.; *n*=11, blue). (**G**) Left, schematic of reciprocal social interaction test following injection of vehicle or captopril (30 mg/kg, i.p.). Right, total social interaction time after captopril (*n*=18, blue) or vehicle (*n*=18, grey). (**H-K**) Time spent huddling (H), interacting nose-to-nose (I), socially exploring (J), or following (K) the partner mouse throughout the assay. (**L**) Diagram of proposed mechanism by which captopril regulates glutamate release onto D1-MSNs via endogenous MERF. (**M**) Central ACE inhibition may provide therapeutic benefit and restore circuit equilibrium in brain conditions characterized by excessive activity of D1-MSNs relative to D2-MSNs. Data are mean ± s.e.m. for all panels; open and closed circles indicate female and male mice, respectively. \**P*<0.05, \*\**P*<0.01, \*\*\**P*<0.001, \*\*\*\**P*<0.0001, ANOVA simple effect of session/treatment (B), and ANOVA treatment main effect (C, G-J); see Table S1 for complete statistics.

To translate an increasingly precise understanding of neural circuit function into therapeutic advances, one strategy is to target molecules known to have enriched expression in specific circuit elements (*1*). Expression of ACE in the striatonigral pathway of the brain was identified decades ago (*5*), but the function of ACE within brain circuits has remained enigmatic. Our data suggest it is not a canonical function related to angiotensin signaling. Instead, we find that ACE regulates endogenous opioid signaling in the NAc by degrading MERF. This action may involve the catalytic N-domain of ACE, which is not essential for regulating blood pressure (*40*) but degrades MERF very efficiently (*31*). Through this mechanism, ACE normally constrains activation of presynaptic mu opioid receptors on glutamatergic axon terminals (Fig. 4, L and M). Pharmacological inhibition of ACE prevents degradation of MERF, thereby enhancing endogenous mu opioid signaling in the NAc. This enhancement of endogenous signaling is similar to the effects of selective reuptake inhibitors for other neurotransmitters, which have substantial therapeutic value for brain disorders. Enhanced endogenous opioid signaling may explain clinical reports that patients taking ACE inhibitors like captopril experience relief from depression and improved quality of life (*6-13*). Captopril has high affinity for the catalytic site located in the ACE N-domain (*41*), and newer ACE inhibitors may lack antidepressant potential because of increased C-domain selectivity or reduced central activity. Chemical optimization of these properties could thus yield new ACE inhibitors specially designed to fine-tune endogenous opioid signaling in the brain.

With growing interest in opioid receptor stimulation as a mechanism of antidepressant action, our data highlight the prospect that enhancement of endogenous opioid signaling may not recapitulate the abuse liability of exogenous opioid agonists. For example, captopril has been prescribed to patients for decades, with no evidence of abuse or dependence. Our data show that systemic captopril administration is not inherently rewarding or aversive in an unbiased place conditioning assay, even though it engages mu opioid receptor signaling. In fact, systemic captopril administration leads to a significant reduction in the rewarding effects of fentanyl, a potent synthetic opioid responsible for many deaths by overdose. This effect can be explained by presynaptic captopril-LTD of excitatory input to D1-MSNs in the NAc (fig. S9), diminishing the activity and ability of this cell type to promote drug reward. Our data suggest that central ACE inhibition has therapeutic potential for a variety of brain conditions characterized by excessive activity of D1-MSNs relative to D2-MSNs. Our findings may thus herald a new era of repositioning and redesigning ACE inhibitors with central activity as a brain circuit-specific pharmacotherapy, emboldened by a long history of safe clinical use.

## Acknowledgements

We thank Drs. Hong Lu and Alan Daugherty (University of Kentucky) for generously providing the floxed ACE mouse line, and the University of Minnesota Mouse Behavior Core for use of facilities to conduct behavioral tests. Mass spectrometry was carried out in the Analytical Biochemistry Shared Resource of the Masonic Cancer Center, supported in part by the U.S. National Institutes of Health and National Cancer Institute (Cancer Center Support Grant CA-77598). Schematics were created with BioRender.com.

## Funding

National Institutes of Health grant R01DA048946 (PER)

National Institutes of Health grant R21 DA050120 (PER)

National Institutes of Health grant F30 DA049476 (BHT)

National Institutes of Health grant F31 MH122094 (CT)

National Institutes of Health grant F30 DA052109 (DDB)

University of Minnesota’s MnDRIVE (Minnesota’s Discovery, Research, and Innovation Economy) initiative (BHT, PER)

## Competing interests

Authors declare that they have no competing interests.

## SUPPLEMENTARY MATERIAL

### Materials and Methods

#### Subjects

All experiments were performed using comparable numbers of both female and male mice (*42*), at 5 to 12 weeks of age. For electrophysiology experiments, we crossed two BAC transgenic reporter lines: Drd1-tdTomato (*43*) (The Jackson Laboratory, stock #016204) and Drd2-eGFP (*44*) (Mutant Mouse Resource & Research Centers #036931-UCD). Hemizygous offspring used for experiments expressed tdTomato in D1-MSNs and/or eGFP in D2-MSNs (Fig. 1C). Floxed *Ace* mice (*45*) (Mutant Mouse Resource & Research Centers #043853-UNC) were crossed with Drd1-Cre BAC transgenic mice (*46*) (FK150; Mutant Mouse Resource & Research Centers #029178-UCD). To generate Ace^fl/fl^ and Ace^wt/wt^ littermates carrying Drd1-Cre for these experiments (Fig. 3, E and F; fig. S6), we crossed parents that were Ace^fl/wt^, with one parent hemizygous for Drd1-Cre. Constitutive *Oprm1* knockout mice (*47*) (The Jackson Laboratory, stock #007559) were crossed with the Drd1-tdTomato reporter line. To generate Oprm1^-/-^ and Oprm1^+/+^ littermates for these experiments (Fig. 3, H to J; fig. S7, E to I), we crossed parents that were Oprm1^+/-^, with one parent hemizygous for Drd1-tdTomato. Constitutive *Penk* knockout mice (*48*) (The Jackson Laboratory, stock #002880) were generated by crossing Penk^+/-^ parents (fig. S2). All genetically modified strains were maintained on a C57BL/6J genetic background, and wildtype C57BL/6J mice were used for LC-MS/MS and behavior experiments. Experimental procedures were conducted between 0900h – 1700h. Mice were housed in groups of 2-5 per cage, on a 12 hour light cycle (0600h – 1800h) at ∼23ºC with food and water provided ad libitum. All procedures conformed to the National Institutes of Health Guidelines for the Care and Use of Laboratory Animals, and were approved by the Institutional Animal Care and Use Committee at the University of Minnesota.

#### Electrophysiology

Mice were deeply anesthetized with isoflurane and perfused with 10 mL ice-cold sucrose cutting solution containing (in mM): 228 sucrose, 26 NaHCO_3_, 11 glucose, 2.5 KCl, 1 NaH_2_PO_4_-H_2_O, 7 MgSO_4_-7H_2_O, 0.5 CaCl_2_-2H_2_O. Mice were subsequently decapitated and brains were quickly removed then placed in ice-cold sucrose cutting solution. Coronal slices (240 µm thick) containing nucleus accumbens (NAc) were collected using a vibratome (Leica VT1000S) and allowed to recover submerged in a holding chamber with artificial cerebral spinal fluid (aCSF) containing (in mM): 119 NaCl, 26 NaHCO_3_, 11 glucose, 2.5 KCl, 1 NaH_2_PO_4_-H_2_O,2.5 CaCl_2_-2H_2_O, 1.3 MgSO_4_-7H_2_O. Slices recovered in warm aCSF (33ºC) for 15 min and then equilibrated to room temperature for at least one hour before use. Slices were transferred to a submerged recording chamber and continuously perfused with aCSF at a rate of 2 mL/min at room temperature. All solutions were continuously oxygenated (95% O2/5% CO_2_).

Whole-cell voltage clamp recordings from MSNs in the NAc core were obtained under visual control using IR-DIC optics from an Olympus BX51W1 microscope. D1-MSNs were distinguished from D2-MSNs by the expression of tdTomato or eGFP, respectively. A subset of experiments (Fig. 3, E and F) used eYFP expression to distinguish D1-MSNs following transfection by AAVDJ-EF1a-DIO-eYFP. Cells were held at -70 mV using borosilicate glass electrodes (3-5 MΩ) filled with (in mM): 120 CsMeSO_4_, 15 CsCl, 10 TEA-Cl, 8 NaCl, 10 HEPES, 5 QX-314, 4 ATP-Mg, 1 EGTA, 0.3 GTP-Na (pH 7.2-7.3). Recordings were performed using a MultiClamp 700B (Molecular Devices), filtered at 2 kHz, and digitized at 10 kHz. Data acquisition and analysis were performed online using Axograph software. Series resistance was monitored continuously and experiments were discarded if resistance changed by >20%.

Excitatory synaptic transmission was pharmacologically isolated using GABAA receptor antagonist picrotoxin (50 µM, Tocris). Excitatory postsynaptic currents (EPSC) were electrically evoked locally using bipolar stimulating electrodes (ISO-flex, AMPI) at 0.05 Hz. Miniature EPSCs (mEPSC) were obtained in the presence of tetrodotoxin (500 nM, Fischer Scientific) to block spontaneous activity. At least 200 events per treatment (e.g. “baseline”) was acquired across 18 sec sweeps, filtered at 0.5 kHz, and detected using an amplitude threshold of 6 pA and a signal-to-noise ratio threshold of 4 standard deviations.

#### Drugs

Electrophysiology experiments used: captopril (10 µM (*18*), Cayman Chemical), Met-enkephalin-Arg-Phe(“MERF” 0.01 µM – 10 µM, Bachem), Met-enkephalin acetate salt (0.01 µM – 10 µM, Bachem), valsartan (2 µM, BioVision), angiotensin I trifluoroacetate salt (1 µM, Bachem), naloxone hydrochloride dihydrate (10 µM(*19*), Millipore Sigma), bestatin (10 µM (*18*), Millipore Sigma), DL-thiorphan (1 µM (*18*), Millipore Sigma),SDM25N hydrochloride (500 nM (*22*), Tocris), nor-Binaltorphimine dihydrochloride (“nor-BNI” 100 nM (*49*), Tocris), CTAP (1 µM (*50*), Tocris). Standards for LC-MS/MS were purchased from Bachem and included: Metenkephalin acetate salt, Leu-enkephalin acetate salt, Met-enkephalin-Arg-Phe, Met-enkephalin-Arg-Gly-Leu, Dynorphin A (1-8) acetate salt, Dynorphin B trifluoroacetate salt, Substance P acetate salt, Angiotensin II acetate salt, and Oxytocin acetate salt. Behavioral experiments used captopril (30 mg/kg i.p.), fentanyl (0.04 mg/kg s.c., Fagron), and sterile saline (5 mL/kg i.p. or s.c.).

#### Liquid Chromatography-Tandem Mass Spectrometry (LC-MS/MS)

Coronal slices (300 µm thick) were individually submerged in 100 µL aCSF (± KCl, ± peptidase inhibitors) and extracellular fluid was collected after 20 minutes and stored at -80ºC. Samples underwent a modified desalting protocol (*51*) with C18-material stage tips (SP301, Thermo Scientific), was eluted with 200 µL solvent (40:60:0.1% water:acetonitrile:trifluoroacetic acid) and dried via speed vacuum overnight. Samples were reconstituted in 12 µL solvent (98:2:0.1% water:acetonitrile:formic acid).

Desalted concentrated samples underwent targeted proteomic identification and quantification based on selected reaction monitoring (SRM) and liquid chromatography-tandem mass spectrometry (*52*). Samples (5 µL) were injected onto a home-packaged analytical C18 reverse phase column (Phenomenex, Torrance, CA) and subsequently eluted with buffer A (0.1% FA in water) and buffer B (0.1% FA in ACN) with the following gradient profile: 0 – 5 min 2% buffer B flow rate at 1 µL/min; 5 – 5.5 min 2% B at 1 – 0.3 µL/min; 5.5 – 25 min 2 – 35% B at 0.3 µL/min; 25 – 26 min 35 – 90% B at 0.3 – 1 µL/min; 26 –29 min 90% B at 1 µL/min; 29 – 30 min 90 – 2% B at 1 µL/min; 30 – 35 min 2% B at 1 µL/min. Mass spectrometry detection was obtained on a TSQ Quantiva Triple Quadrupole (Thermo Scientific) in positive nanospray ionization mode. Mass spectrometry conditions were: spray voltage 2.0 kV, ion transfer tube temperature 350 °C, collision energy 4 – 29.2 V, and collision gas (argon) pressure 1 mTorr. Resolution settings were 0.7 Da (full width at half-maximum) for both quadrupoles and transition dwell times were 15 ms. Standards for absolute peptide quantification (10 pm, 50 pm, 100 pm, 500 pm, 1 nm, 5 nm, 10 nm) were injected after experimental samples and contained: Met-enkephalin, Leu-enkephalin, Met-enkephalin-Arg-Phe (MERF), Met-enkephalin-Arg-Gly-Leu (MERGL), Dynorphin A (1-8), Dynorphin B, Substance P, Angiotensin II, and Oxytocin. Although MERGL and oxytocin were included in the standards and detected reliably with comparable sensitivity (Fig. S1), these peptides were not detected in experimental samples.

Skyline (MacCoss Lab) was used to empirically determine SRM transitions for all peptide standards and quantitative data processing. SRM transitions corresponding to the five largest integrated peaks were selected for targeted proteomic analysis and precursor / product transitions with the largest peak area was used for absolute peptide quantification as derived from calibration curves: Met-enkephalin (574.2330 / 278.1135), Leu-enkephalin (556.2766 / 278.1135), MERF (439.2049 / 714.3392), MERGL (450.7235 / 737.3763), Dynorphin A (1-8) (327.8591 / 409.7534), Dynorphin B (524.2999 / 627.3487), Substance P (674.3713 / 600.3378), Angiotensin II (523.7745 / 263.1390), Oxytocin (504.3700 / 495.8460). Peaks were manually inspected to ensure correct detection and integration for each peptide per sample.

#### Behavior

##### Place Conditioning

As previously described (*53*), activity test chambers (Med Associates, 11 × 11 × 11 in.) consisted of a single compartment with two types of removable floor tiles: transparent plastic with a rough texture and black plastic with a smooth texture. A five-day protocol was used which included a 20 minute baseline session on day 1 that included both contexts; three conditioning days consisting of one 30 minute saline (5 mL/kg s.c.) conditioning session in the morning (AM) paired to one context and one 30 minute vehicle + fentanyl (0.04 mg/kg s.c.), captopril (30 mg/kg i.p.) + fentanyl, captopril, or vehicle conditioning session paired to the other context in the afternoon; and finally a 20 minute test session on day 5 containing both contexts. Conditioning sessions were at least three hours apart. Captopril or vehicle was given 30 minutes and fentanyl was given 5 minutes prior to placement in activity chambers on conditioning days. Wildtype C57BL/6J mice were tested at 6 to 10 weeks of age and randomly assigned to either the control, captopril + fentanyl, or captopril group. All mice habituated to the procedure room for one hour prior to placement in activity chambers. Based on baseline preference as measured by time spent in either contexts, animals were assigned to either a smooth black or rough transparent context for saline and the opposite context for vehicle + fentanyl, captopril + fentanyl, captopril, or vehicle so that the group average baseline preference was approximately 50%. Individuals with a baseline preference greater than 75% for any one context was excluded. The total ambulatory distance in each activity chamber was recorded with Med Associates software during all tests and conditioning sessions.

##### Social Interaction Test

Evaluation of social interaction was performed as previously described (*54*). All experiments were conducted at 60-70 luminosity, and at temperature conditions equal to those of the animal housing facility. Experimental sessions were video recorded and social interaction was hand scored by researchers blind to experimental conditions. Wildtype C57BL/6J mice were moved to an isolated testing room 1 hour before tests. Mice were tested at 6 to 8 weeks of age in an opaque white rectangular box with 1 cm of fresh corn cob bedding on the floor for 10 minutes. Both mice were novel social partners, age- and sex-matched, were not siblings or cage mates, and were both given either captopril (30 mg/kg i.p.) or vehicle. Video recordings of social behaviors exhibited by mice were hand scored by a blinded experimenter using Button Box 5.0 (Behavioral Research Solutions, LLC). Social behaviors were categorized into one of the following groups (*55*): nose-nose interaction (direct investigation of orofacial region), huddling (stationary sitting next to partner), social exploration (anogenital investigation, social sniffing outside of orofacial region, social grooming), and following. The sum of these social behaviors were used for total “Interaction Time” (Fig. 4G).

#### RT-qPCR

Quantitative RT-PCR was performed on tissue punches containing the dorsal and ventral striatum, as previously described (*56*). Tissue was snap frozen on dry ice and stored at -80°C. RNA was extracted and isolated using the RNeasy Mini Kit (Qiagen) according to manufacturer instructions. A NanoDrop One microvolume spectrophotometer (Thermo Fisher Scientific, Waltham, MA) was used to measure RNA concentration and verify that samples had A260/A280 purity ratio ≥ 2. Reverse transcription was performed using Superscript III (Invitrogen, Eugene, OR). For each sample, duplicate cDNA reactions and subsequent qPCR reactions were conducted in tandem on both samples. Mouse β-actin mRNA was used as the endogenous control to measure differences in expression of ACE mRNA. Primer sequences for detection of ACE mRNA were forward 5’-GCCTCCCAAGGAATTAGAAGAG-3’ and reverse 5’-TGATGTACTCGGACCCATAGT-3’. Quantitative RT-PCR using SYBR green (BioRad, Hercules, CA) was carried out with a Lightcycler 480 II (Roche, Basel, Switzerland) using the following cycle parameters: 1 x (30 sec @ 95°C), 35 x (5 sec @ 95°C followed by 30 sec @ 60°C). Results were analyzed by comparing the C(t) values of the treatments tested using the ΔΔC(t) method. Expression values of target genes were first normalized to the expression value of β-actin. The mean of cDNA replicate reactions was used to quantify the relative target gene expression.

#### Statistical Analysis and Data Presentation

Female and male mice were used in all experiments, and sex was included as a variable in factorial ANOVA models analyzed using IBM SPSS Statistics v24 or GraphPad Prism 8. Significant interactions provided strong evidence that the effect of one variable depended on the level of another variable (*57*), and were decomposed by analyzing simple effects (i.e., the effect of one variable at each level of the other variable). Significant main effects were further analyzed using Fisher’s LSD post-hoc tests. The Type I error rate was set to α=0.05 (two-tailed) for all comparisons. In the figures, significant ANOVA effects and follow-up tests are denoted by black asterisks that indicate \**P*<0.05, \*\**P*<0.01, \*\*\**P*<0.001, \*\*\*\**P*<0.0001. All data are presented mean ± s.e.m., with open and closed circles indicating data points from female and male mice, respectively. Complete statistical information can be found in Table S1.

**Fig. S1.**
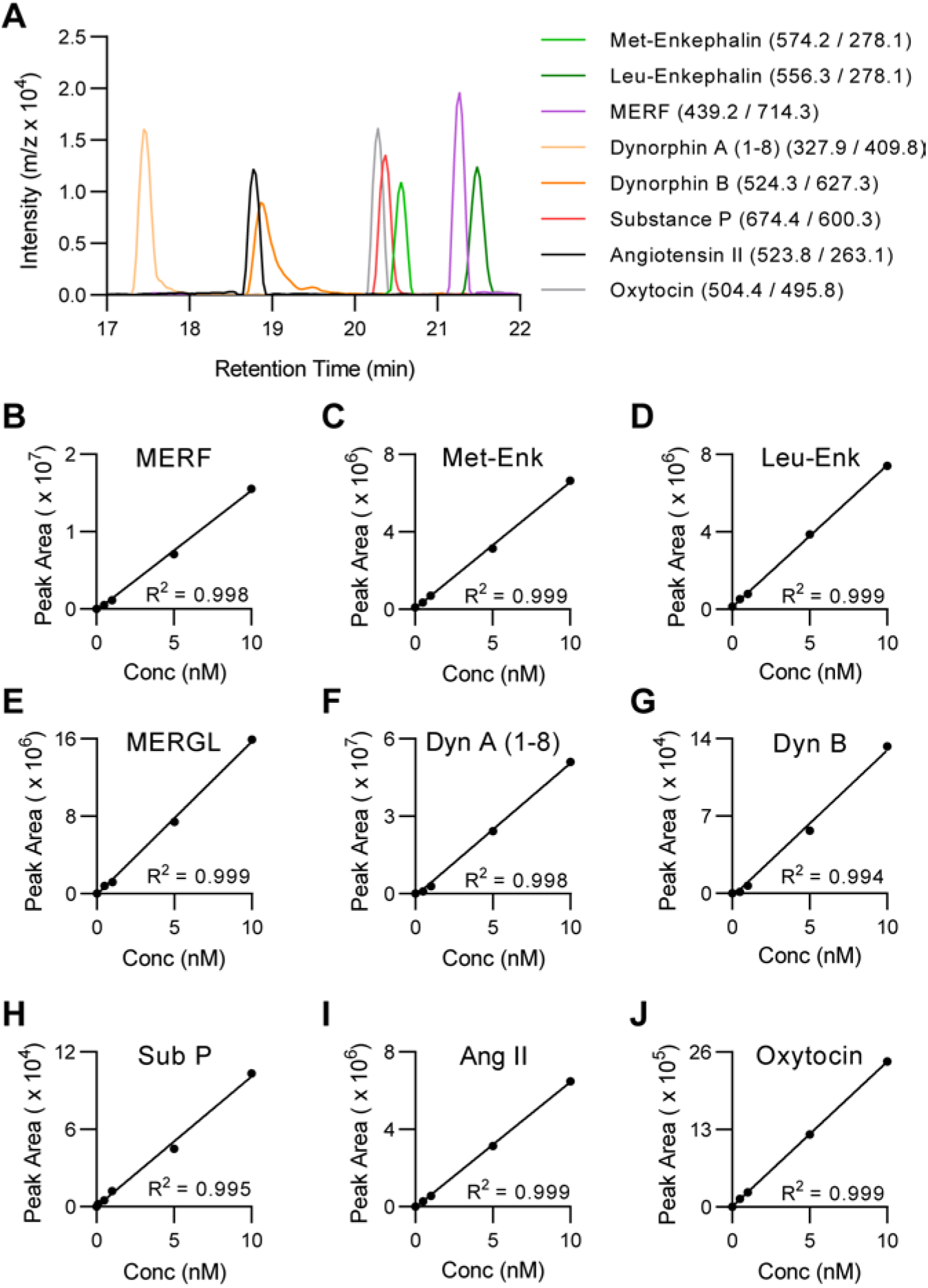
Liquid chromatography-tandem mass spectrometry can be used for targeted neuropeptide identification and quantification. (**A**) Reconstructed chromatogram showing typical retention time and SRM transition (precursor / product) that produces the largest integrated peak area (out of five different transitions per peptide, not shown) which is used for peptide quantification. (**B-J**) Calibration curves for each peptide (10 pM, 50 pM, 100 pM, 500 pM, 1 nM, 5 nM, and 10 nM, black dots) from standards containing mixture of all peptides.

**Fig. S2.**
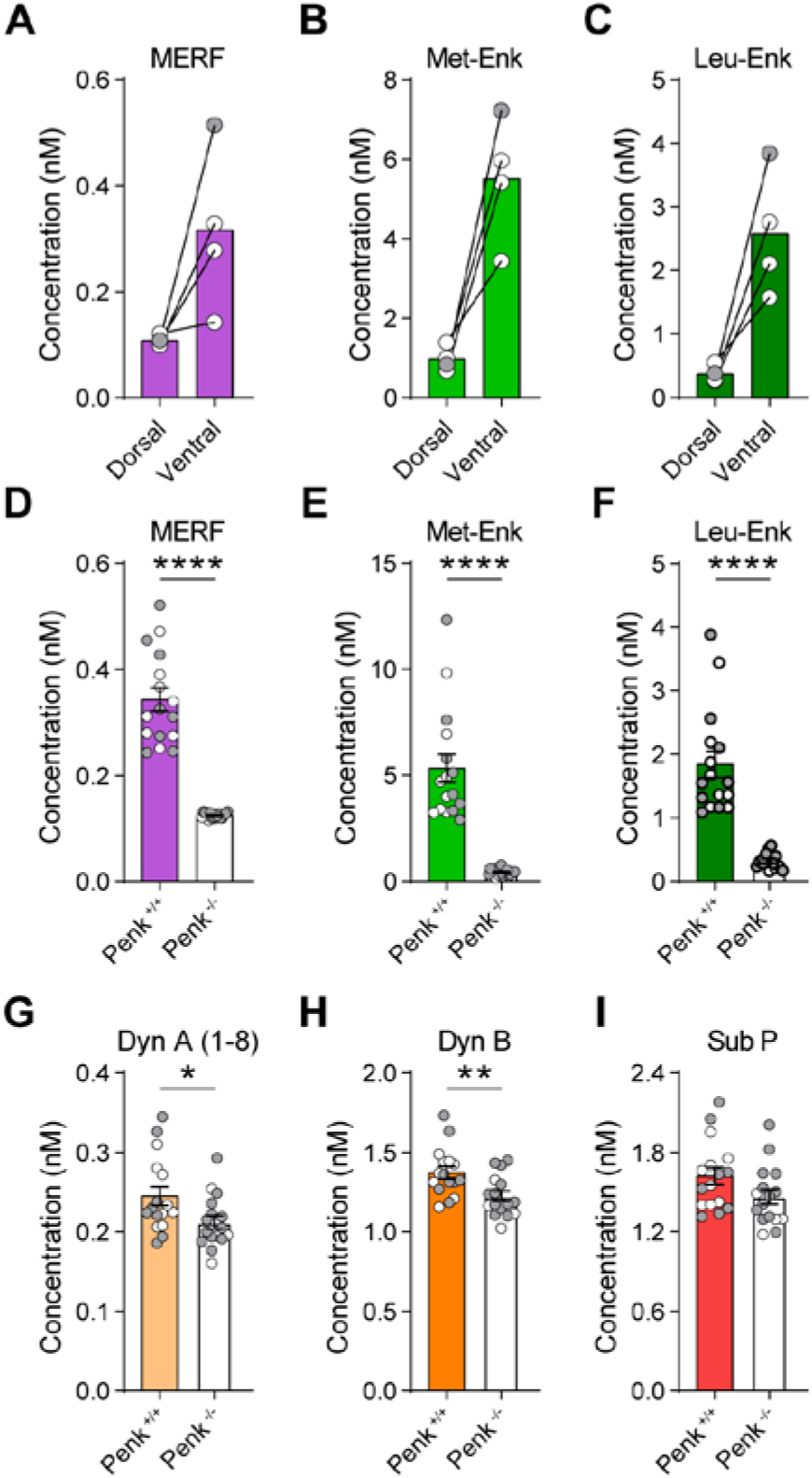
Neuropeptide quantification by LC-MS/MS is sensitive to regional and genetic effects. (**A-C**) Extracellular levels of MERF, Met-, and Leu-enkephalin from ventral striatum (nucleus accumbens) and dorsal striatum tissue punches after submersion in aCSF with KCl (50 mM, n=4). (**D-I**) Quantification of peptides after KCl stimulation of individual striatal slices from constitutive Penk knockout mice (Penk^-/-^, n=16, white bars) compared to wildtype littermates (Penk^+/+^, n=16, color bars) to validate specificity and sensitivity of SRM-based targeted proteomic quantification for endogenous enkephalin peptides. Data are mean ± s.e.m. for all panels; open and closed circles indicate samples from female and male mice, respectively. *P<0.05, **P<0.01, ***P<0.001,****P<0.0001, two-sample t-test (D-I); see Table S1 for complete statistics.

**Fig. S3.**
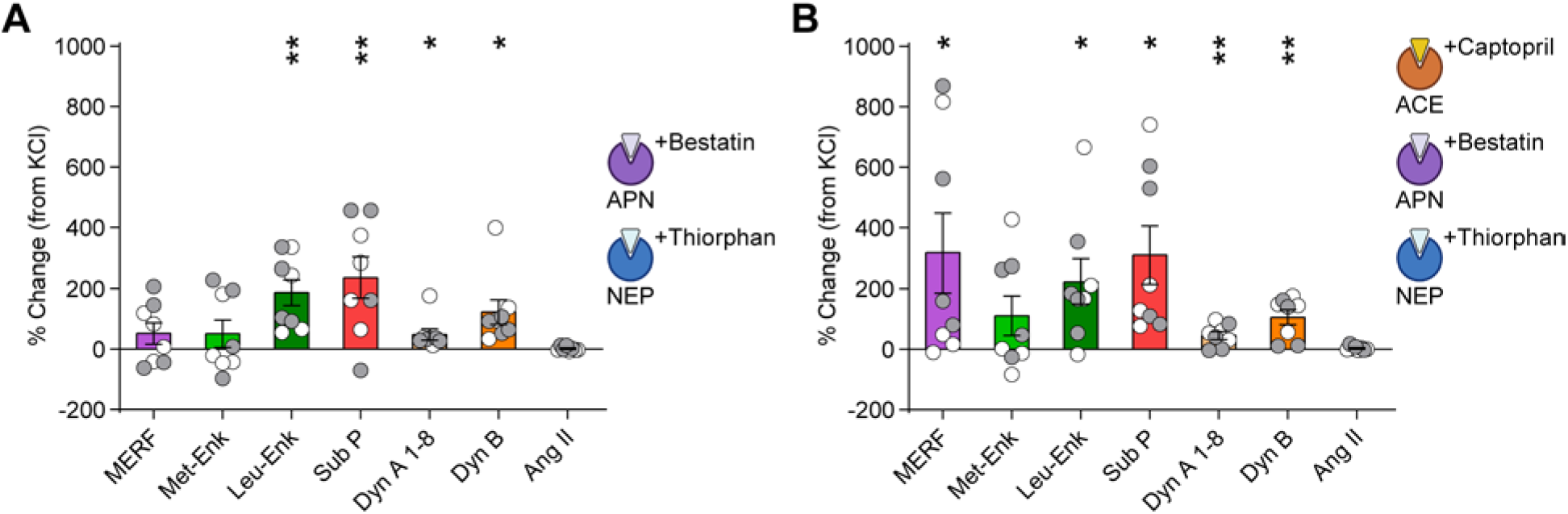
Double dissociation between regulation of MERF and conventional enkephalins. (**A**) Quantification of neuropeptides from striatal slices (*n*=8) following submersion in aCSF with KCl (50 mM) plus peptidase inhibitors bestatin (10 µM) and thiorphan (1 µM). (**B**) Neuropeptide levels of KCl-stimulated slices after exposure to a cocktail of captopril (10 µM), bestatin, and thiorphan reflect additive effects from samples exposed to captopril alone seen in Figure 2c and combined bestatin plus thiorphan seen in (A). Data are mean ± s.e.m. for all panels; open and closed circles indicate samples from female and male mice, respectively. \**P*<0.05, \*\**P*<0.01,\*\*\**P*<0.001, \*\*\*\**P*<0.0001, one-sample *t*-test; see Table S1 for complete statistics.

**Fig. S4.**
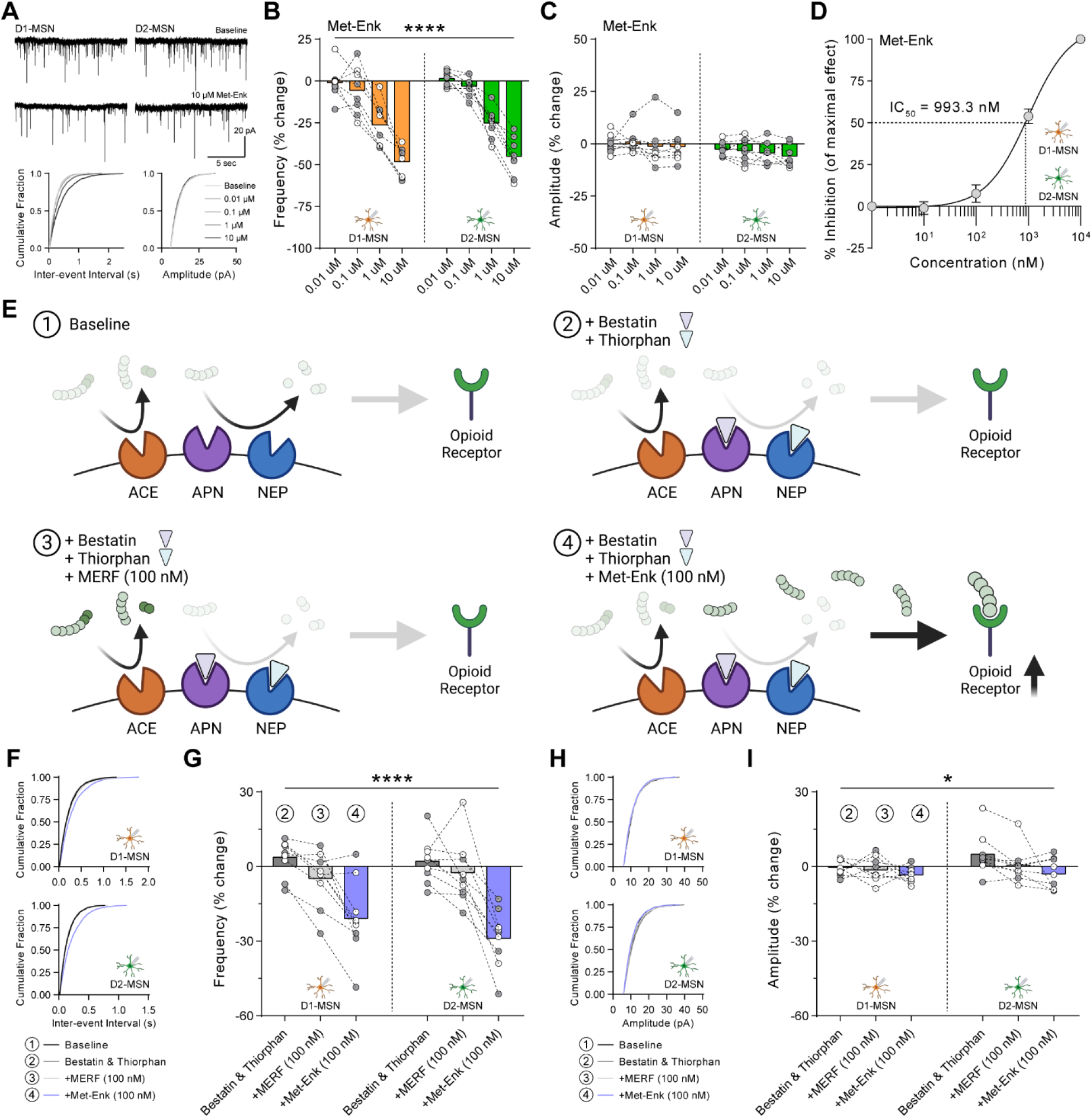
Met-enkephalin synergizes with peptidase inhibitors of aminopeptidase N and neprilysin to depress glutamate release. (**A**) Top, representative traces of mEPSCs from D1-MSNs (left) and D2-MSNs (right) during baseline and after bath perfusion of Met-enkephalin (10 µM). Bottom, cumulative fraction plots of inter-event interval (left) and amplitude (right) of mEPSC events at increasing Met-enkephalin concentrations (0.01 µM – 10 µM). (**B, C**) Met-enkephalin caused a dose-dependent decrease in mEPSC frequency (B) and did not affect mEPSC amplitude (C) in D1-MSNs (left, orange, *n*=8) and D2-MSNs (right, green, *n*=8). (**D**) Sigmoidal interpolation of Met-enkephalin dose-response normalized to maximal frequency change at 10 µM (IC_50_: 993.3nM, 95% CI: 669.1 – 1475 nM, *n*=16). (**E**) Conceptual schematic showing how peptidase inhibitors bestatin and thiorphan toward aminopeptidase N and neprilysin, respectively, affect opioid receptor signaling by MERF or Met-enkephalin. (**F-I**) Effect of combined bestatin and thiorphan plus MERF or Met-enkephalin in recordings of D1-MSNs (*n*=9) or D2-MSNs (*n*=9) on mEPSC cumulative fraction plots of inter-event interval (F), percent change in frequency (G) cumulative fraction plots of amplitude (H), and percent change in amplitude (I). Data are mean ± s.e.m. for all panels; open and closed circles indicate recordings from female and male mice, respectively. \**P*<0.05, \*\**P*<0.01, \*\*\**P*<0.001, \*\*\*\**P*<0.0001, ANOVA concentration or treatment main effect (B, G, I); see Table S1 for complete statistics.

**Fig. S5.**
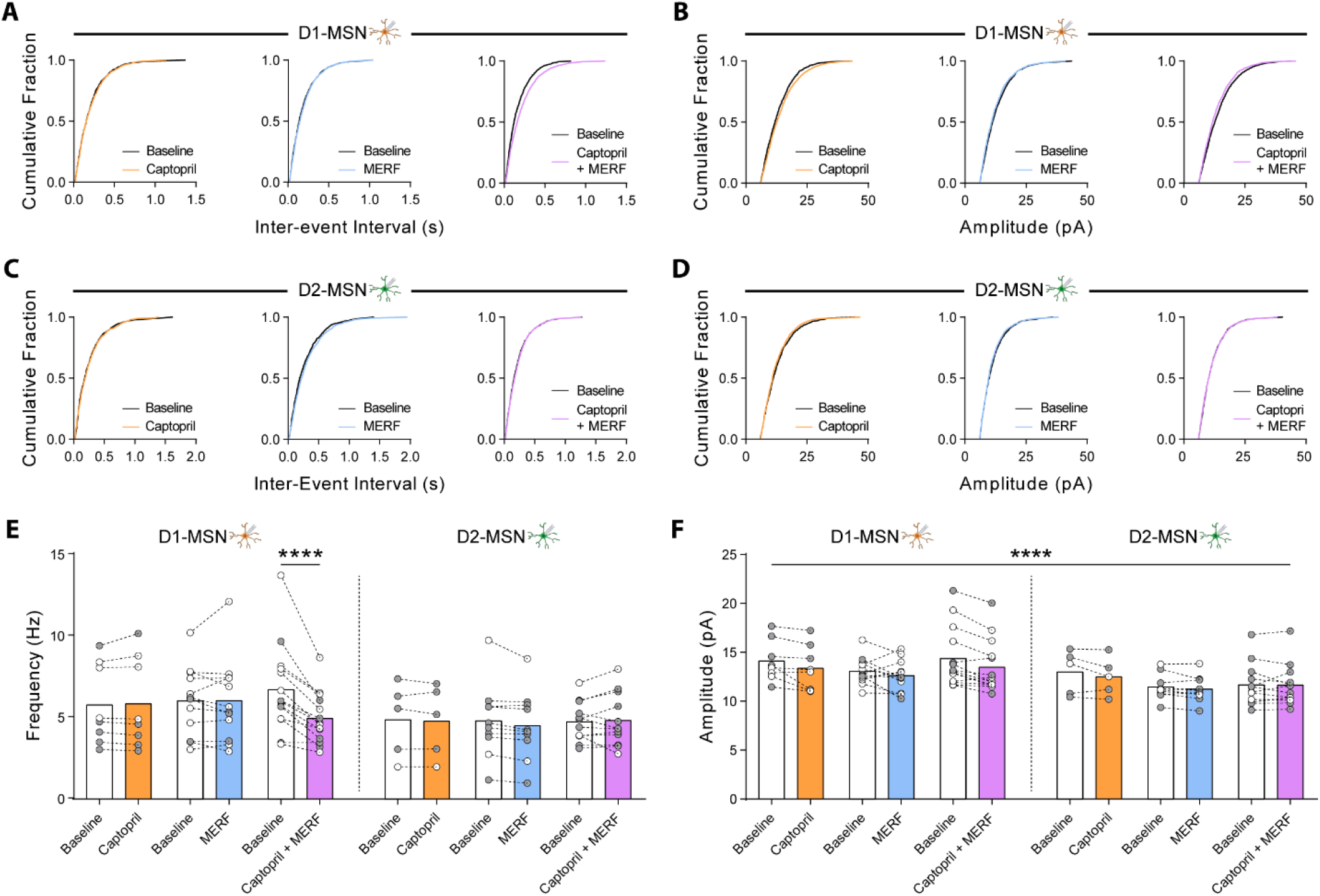
MERF and captopril synergistically depress presynaptic glutamate release. (**A, B**) Cumulative fraction plots of mEPSC inter-event interval (A) and amplitude (B) in D1-MSNs before and after captopril (10 uM, *n*=8), MERF (100 nM, *n*=12), or combined (*n*=14). (**C, D**) Cumulative fraction plots of mEPSC inter-event interval (C) and amplitude (D) in D2-MSNs before and after captopril (*n*=5), MERF, (*n*=10), or combined (*n*=12). (**E, F**) Average mEPSC frequency (E) and amplitude (F) before and after captopril, MERF, or combined in D1-MSNs (left) and D2-MSNs (right). Data are mean ± s.e.m. for all panels; open and closed circles indicate recordings from female and male mice, respectively. \**P*<0.05, \*\**P*<0.01, \*\*\**P*<0.001, \*\*\*\**P*<0.0001, ANOVA followed by Fisher’s LSD post-hoc test (E) and ANOVA time main effect (F); see Table S1 for complete statistics.

**Fig. S6.**
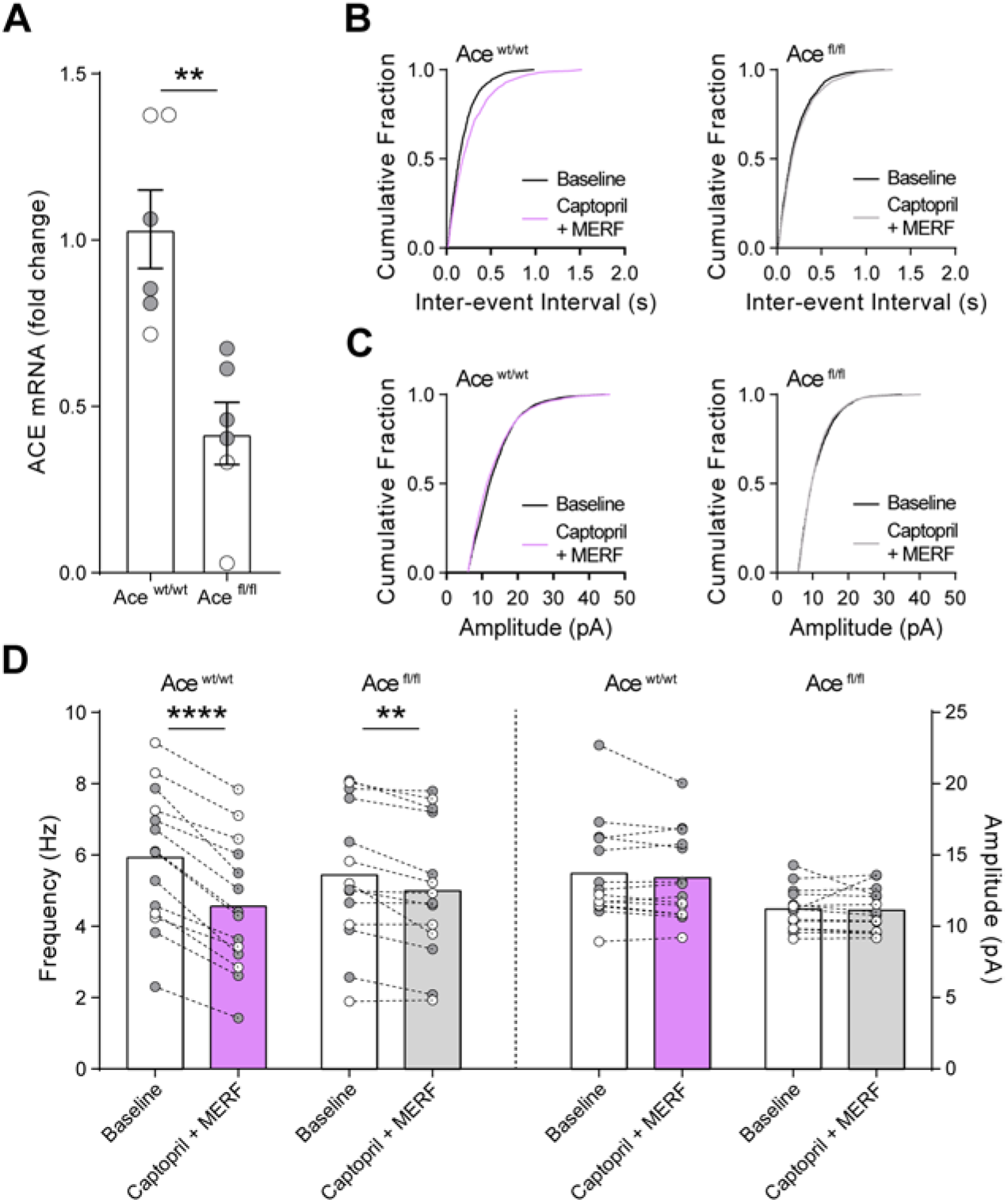
ACE is necessary for synergistic effect of captopril and MERF. (**A**) mRNA quantification from striatal tissue punches in control (Ace^wt/wt^; Drd1-Cre, *n*=6) and conditional ACE knockout mice (Ace^fl/fl^; Drd1-Cre, *n*=6). (**B**-**D**) mEPSC parameters before and after combined captopril (10 uM) and MERF (100 nM) in Ace^wt/wt^ (*n*=14) and Ace^fl/fl^ (*n*=14) mice shown as cumulative fraction plots of inter-event interval (B), cumulative fraction plots of amplitude (C), and average frequency (left) and amplitude (right, D). Data are mean ± s.e.m. for all panels; open and closed circles indicate female and male mice, respectively. \**P*<0.05, \*\**P*<0.01, \*\*\**P*<0.001,\*\*\*\**P*<0.0001, ANOVA genotype main effect (A) and ANOVA followed by Fisher’s LSD post-hoc test (D); see Table S1 for complete statistics.

**Fig. S7.**
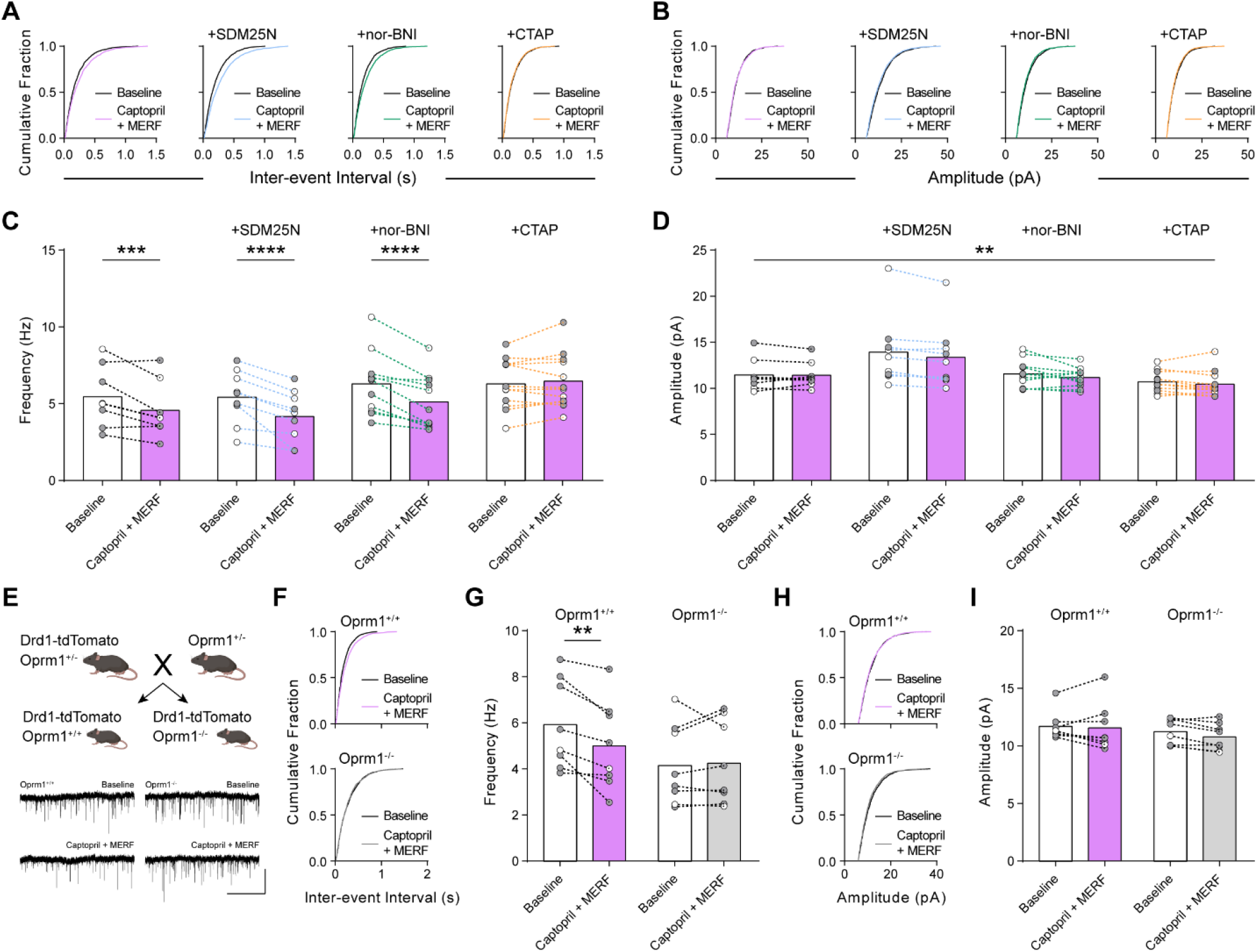
Mu opioid receptor signaling is necessary to reduce glutamate release by combined captopril and MERF. (**A-D**) mEPSC parameters shown as cumulative fraction plots of inter-event interval (A), amplitude (B), average frequency (C), or average amplitude (D) before and after combined (*n*=8) captopril (10 uM) and MERF (100 nM) plus selective antagonists of delta (SDM25N, 0.5 µM, blue, *n*=9), kappa (nor-BNI, 0.1 µM, green, *n*=11), or mu (CTAP, 1 µM, orange, *n*=12) opioid receptors. (**E**) Top, genetic cross to obtain control (Oprm1^+/+^; Drd1-tdTomato) and constitutive mu opioid receptor knockout (Oprm1^-/-^; Drd1-tdTomato) mice. Bottom, representative traces of mEPSCs in D1-MSNs from Oprm1^+/+^ (left) and Oprm1^-/-^ (right) mice during baseline and after combined captopril and MERF. (**F-I**) mEPSC parameters shown as cumulative fraction plots of inter-event interval (F), amplitude (H), average frequency (G), or average amplitude (I) before and after combined captopril and MERF in Oprm1^+/+^ (*n*=8) and Oprm1^-/-^ (*n*=8) mice. Data are mean ± s.e.m. for all panels; open and closed circles indicate recordings from female and male mice, respectively. \**P*<0.05, \*\**P*<0.01, \*\*\**P*<0.001,\*\*\*\**P*<0.0001, ANOVA followed by Fisher’s LSD post-hoc test (C, G) and ANVOA time main effect; see Table S1 for complete statistics.

**Fig. S8.**
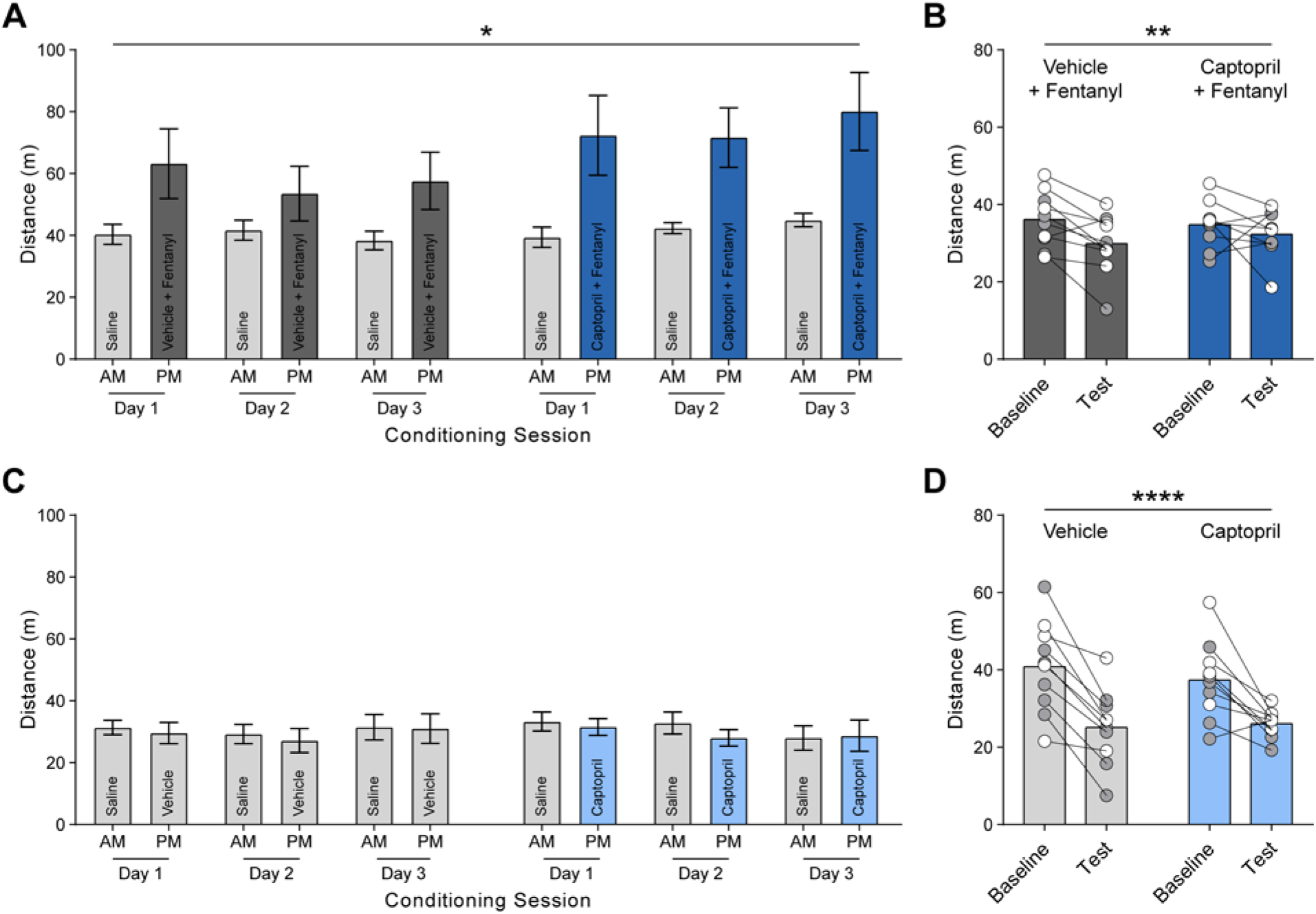
Captopril does not alter locomotion. (**A, B**) Total ambulatory distance of mice receiving vehicle plus fentanyl (0.04 mg/kg, s.c.; *n*=10) or captopril (30 mg/kg, i.p.) plus fentanyl (*n*=9) throughout unbiased place conditioning assay during conditioning sessions (A) and on baseline and test day (B). (**C, D**) Total ambulatory distance of mice receiving vehicle (*n*=10) or captopril (*n*=10) throughout unbiased place conditioning assay during conditioning sessions (C) and on baseline and test day (D). Data are mean ± s.e.m. for all panels; open and closed circles indicate female and male mice, respectively. \**P*<0.05, \*\**P*<0.01, \*\*\**P*<0.001, \*\*\*\**P*<0.0001, ANOVA time main effect (A, B, D); see Table S1 for complete statistics.

**Fig. S9.**
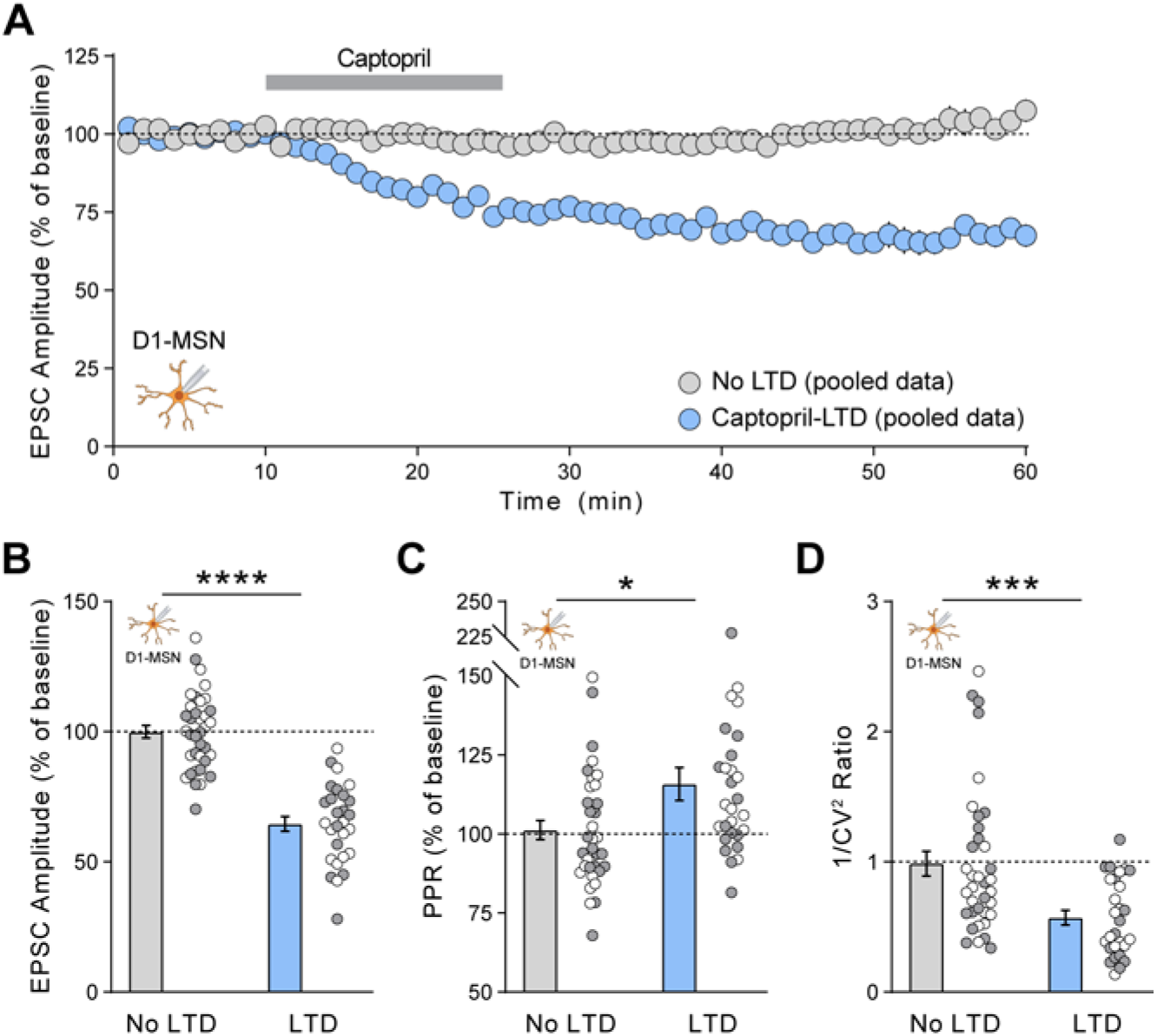
Captopril-LTD in D1-MSNs occurs via a presynaptic mechanism. (**A**) Time course of captopril-LTD from pooled LTD (*n*=28) and control (*n*=36) data sets. (**B-D**) EPSC parameters averaged in the last 5 min of each recording: EPSC amplitude (B), paired-pulse ratio (C), and inverse of squared-coefficient of variation (D). Pooled data includes recordings from D1-MSNs seen in Figure 1 and Figure 3I, J. Data are mean ± s.e.m. for all panels; open and closed circles indicate recordings from female and male mice, respectively. \**P*<0.05, \*\**P*<0.01,\*\*\**P*<0.001, \*\*\*\**P*<0.0001, ANOVA LTD main effect (B, C, D); see Table S1 for complete statistics.

## Notes

### Competing Interest Statement

The authors have declared no competing interest.

